# An eye-tracking study of visual attention in chimpanzees and bonobos when viewing different tool-using techniques

**DOI:** 10.1101/2024.08.01.605994

**Authors:** Yige Piao, James Brooks, Shinya Yamamoto

## Abstract

Chimpanzees and bonobos are excellent tool users and can socially learn various skills. Previous studies on social learning mainly measure success/failure in acquiring new techniques, with less direct measurement of proximate mechanisms like visual attention during the process. This study investigates how apes observe tool-using demonstrations through eye-tracking. After checking initial techniques, six chimpanzees and six bonobos were shown video demonstrations of human demonstrators using a tube to dip (low-efficiency) or suck (high-efficiency) juice, and then tried the task themselves. Attention to each video was compared to participants’ knowledge. Although no individuals acquired the high-efficiency technique through video demonstrations, eye-tracking results revealed attentional differences between individuals familiar with different techniques. Compared with individuals already familiar with both techniques, individuals knowing only the dipping technique showed less attention to the unfamiliar sucking technique. This result indicates that apes may not attend much to what they do not know well, which aligns with reported interplay of action observation and understanding. Attentional patterns to specific areas was similar between species, though there was a tendency towards more attention to faces in bonobos and food in chimpanzees. This study emphasizes the importance of detailed investigation into social learning process using eye-tracking.

## Introduction

Chimpanzees and bonobos, our evolutionary closest relatives, stand out among non-human tool-users in both repertoire and ability to socially learn [1], providing significant insight into the evolution of human culture [2–4]. Wild chimpanzees demonstrate an impressive array of tool-use, including underground termite fishing and nut-cracking [5–8] in their material culture [9–11]. While wild bonobos seldom use tools [12,13], bonobos in captivity are as skilled in tool-use as chimpanzees [14–16]. Though individuals may innovate behaviors through asocial learning [17], social learning is pivotal in the transmission and maintenance of such behaviors [1,18–20] and even leads to local traditions and primitive culture [21–24].

Over the decades, social learning of tool-use has been a hot topic in cognitive studies of great apes [1,25,26]. However, previous studies mostly focus on the success/failure in acquiring new skills and the transmission network [27–30], while the underpinning mechanisms remain underexplored. More specifically, little is known about how observers view demonstrations during this process. Because social learning involves obtaining information from other individuals [31], investigating this information extraction process is essential to fully understand social learning and its role in facilitating traditions and cultures.

Eye-tracking technology provides qualitative and quantitative investigation into the cognitive process of non-human primates [32]. Here we investigate how *Pan* species view demonstrations of familiar and unfamiliar tool-using techniques. We adopted a simple tool-using task used in a previous study [33]. Apes could use a transparent tube to obtain grape juice through a small hole. The tube can be used as either a stick to dip (low-efficiency) or a straw to suck (high-efficiency). Within a group, multiple techniques may be innovated based on individual experience and knowledge, providing the opportunity to investigate how social attention to differing techniques is affected by prior knowledge [34,35].

We first checked participants’ initial tube-using technique, and then used eye-tracking technology to investigate their visual attention when observing human demonstrators displaying the two alternate solutions (which differ in efficiency). We predicted that visual attention would differ both between individuals with differing initial techniques and between bonobos and chimpanzees. More specifically, previous studies indicate a tight link between action observing, understanding and execution [36,37]. Therefore, our first hypothesis was : (1) visual attention will differ between participants with different baseline techniques, especially regarding the more challenging high-efficiency sucking technique. Based on the attentional and motivational differences between the species [38,39], our second hypothesis was: (2) chimpanzees would look more at the action part (i.e. tube and hand), while bonobos would look more at demonstrators’ face. As the key information of the techniques is mainly displayed in the action part, if the second hypothesis was correct, we would expect dipping-technique chimpanzees to be more likely to notice the differences between two techniques. Therefore, the third hypothesis was: (3) dipping-technique chimpanzees would be more likely than bonobos to notice the different technique and switch to the higher-efficiency sucking method.

## Methods

### 1. Animals

Six chimpanzees (*Pan troglodytes verus*) and six bonobos (*Pan paniscus*) at Kumamoto Sanctuary, Japan, participated in this study (see details in supplementary materials).

Participants were first offered several transparent plastic tubes (diameter: inner 4 mm, outer 7 mm; length around 23 cm) and a bottle of grape juice (inner diameter 4 cm, height 19 cm). Tubes could be used either to dip (low-efficiency) or as a straw to suck (high-efficiency) the juice. Participants’ initial method was recorded as their knowledge of tube-using technique.

### 2. Eye-tracking setup

The procedures were similar to previous studies [39,40]. An infrared eye tracker (300 Hz, TX300, Tobii Technology AB) was used to record apes’ visual attention. A 23-inch LCD monitor (43 × 24°, resolution 1280 × 720 pixels) displayed the stimuli from a distance of 60 cm to the participant. During eye-tracking, the participant was provided with grape juice via a custom-made juice dispenser. Automated calibration was conducted for each participant by presenting a video clip on two reference points, and its result was shown on the monitor by presenting small reference icons. This calibration was repeated whenever necessary before each eye-tracking recording. Calibration errors are generally within one degree for most participants [41].

Visual stimuli were two videos in which two human experimenters each displayed either dipping or sucking techniques (Video S1). All participants were familiar with both experimenters for many months to years. Each video lasted 24 seconds, with the first demonstrator (J.B.) performing the action four times in 12 seconds and the second demonstrator (Y.P.) doing the same in another 12 seconds. In dipping-technique videos, demonstrators dipped one end of the tube into the juice through a hole (diameter 1.2 cm) on the panel, retrieved and put that end into the mouth. In sucking-technique videos, demonstrators put one end of the tube into the juice through the hole and sucked some juice via the other end. The juice was completely visible when moving inside the tube, and the liquid level in the bottle was visibly decreasing.

### 3. General procedure

In each trial, participants were called by their names to enter the testing room. Trials began when the participant voluntarily approached the panel and were shown the juice bottle and tube. participants then watched the video demonstrations while their gaze was recorded via eye-tracking. After demonstrations, the experimenter again showed the juice bottle and added some juice into it. Finally, a tube was provided through the hole on the panel, and the juice bottle was attached below this hole using a suction cup (diameter 25 mm). The participant was offered one minute to try the task and her/his behaviors were video recorded. This tube-using task ended either when the participant consumed all juice in the bottle or after one minute had elapsed. If participants left or looked away during the task, every 20 seconds the experimenter attempted to attract their attention by calling the name, jiggling the tube, or adding more juice. Another tube was be provided if the participant left the previous one beyond arm’s reach.

Each participant underwent five trials, with an inter-trial interval of at least five days. Among all trials, dipping- and sucking-technique videos were shown in a counter-balanced order.

### 4. Analysis

Eye movement was filtered using Tobii Fixation Filter with default parameters. Areas of interest (AOI) was defined for each video in the Tobii Studio software (ver. 3.2.1) and included the face (demonstrators’ face), food (juice bottle), action (tube in one hand), and screen (the whole screen) (Figure 1). Statistical analyses were performed in R (v.4.3.2; R Core Team, 2024) using linear mixed effects models (LMM) (‘lmer’ in the package ‘lme4′) with Gaussian error structure and identity link function. The variable Trial was standardized to a mean of 0 and standard deviation of 1 (using the ‘standardize’ function) according to [42]. We used the “check_model” function in the package “performance” to assess different aspects of a model’s fit by visual inspection of the diagnostic plots.

**Figure 1.**
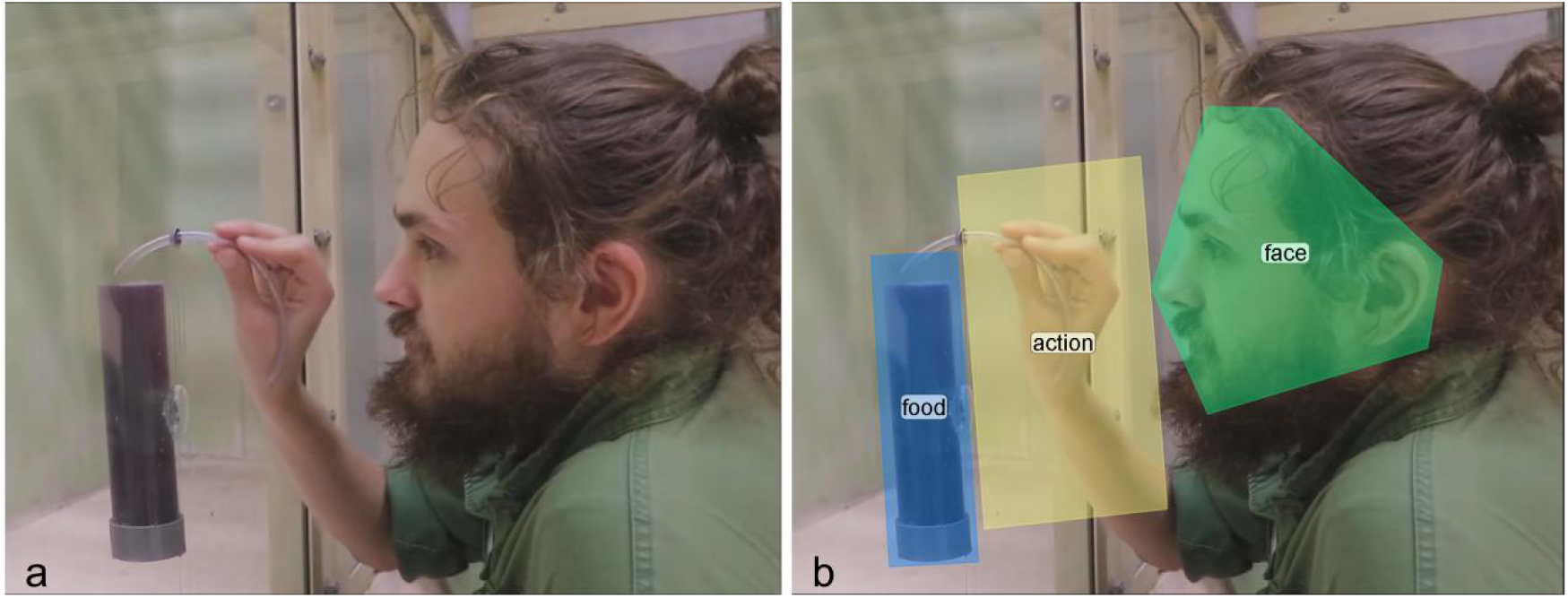
An example of video stimuli. A experimenter (J.B.) demonstrating the tube-using technique (a) and with main AOIs indicated (b).

### Model: looking duration by Knowledge*Stimuli

We first investigated how participants with different tube-using knowledge viewed the videos. Total fixation duration was used for overall attention to the screen, and proportion of looking duration was used for each AOI (e.g. the proportion of looking duration to face is 0.5 if attention to face for 6 seconds with attention to the screen for 12 seconds). Separate models were run for each dependent variable. In all models, we included Trial (trial 1 to 5), Species (bonobos/chimpanzees), Demonstrator (first/second demonstrator), Knowledge (dipping/sucking), Stimuli (dipping/sucking videos), the interaction between Knowledge and Stimuli, and stimulus order (whether the first presented video was dipping/sucking), as fixed effects. We included participant identity (ID) to account for repeated measures for each individual, and the effects of Trial and Stimuli varying by ID, as random effects and random slopes, respectively. The random-effects structure was kept maximal to save conservativity of the tests according to [43]. The model syntax used for all models in main text was: the proportion of Looking duration ∼ Trial + Species + Demonstrator + Knowledge*Stimuli + First-show + (1 + Trial + Stimuli || ID). We confirmed the normality of residuals, homogeneity of variance, and normality of random effects (ID) in all models. We also checked Variance Inflation Factors (VIF) and found that collinearity remained low in all models (VIF < 5). When a significant interaction effect was detected in the model, we further examined it by testing simple effects in the subsets of data including each level of predictors.

As exploratory analyses, we additionally tested whether the behavioral responses of participants might indicate how they observed the stimuli, as well as analyzed two smaller AOIs (eye and mouth), to investigate potential differences in participants’ attention. The modelling approach was similar to the above and results are reported in supplementary materials.

## Results

In total, six chimpanzees participated in 30 eye-tracking tests and 25 tube-using tasks in the testing room. Two individuals, Zamba and Misaki, were often eager to leave the room after eye-tracking and some of their tube-using tasks were therefore conducted in a room adjacent to the outdoor enclosure immediately upon return. All bonobos completed the eye-tracking and tube-using tasks in their testing room. A naive observer coded 30% of the video records of tube-using tasks and showed high inter-rater reliability (Cohen’s Kappa test, Kappa value = 0.912 , z = 6.2, *P* < 0.001) with the experimenter (Y.P.).

### Looking duration by Knowledge*Stimuli

A significant interaction between knowledge and stimuli was detected between knowledge and video stimulus for overall attention to the videos (β = 1.814, SE = 0.657, χ ^2^ = 7.618, *P* = 0.020) (Figure 2). We therefore tested simple effects in the subsets of data on each level of test predictors. Sucking-technique participants looked significantly more at sucking-technique videos than dipping-technique participants (β = 4.245, SE = 1.340, χ ^2^ = 10.037, *P* = 0.012). There was also a marginal effect of sucking-technique participants attending more to dipping-technique videos than dipping-technique participants (β = 2.871, SE = 1.284, χ^2^ = 4.997, *P* = 0.051). Across participants and stimuli, attention to videos decreased from Trial 1 to 5 (β = - 0.696, SE = 0.147, χ^2^ = 22.382, *P* < 0.001). Similarly, participants looked less at the second demonstrator in the video (β = - 1.373, SE = 0.274, χ ^2^ = 25.099, *P* < 0.001). Both species paid similar attention to the screen overall ( χ ^2^ = 0.116, *P* = 0.741), and the order of presenting videos did not influence their looking duration ( χ ^2^ = 1.634, *P* = 0.206).

**Figure 2.**
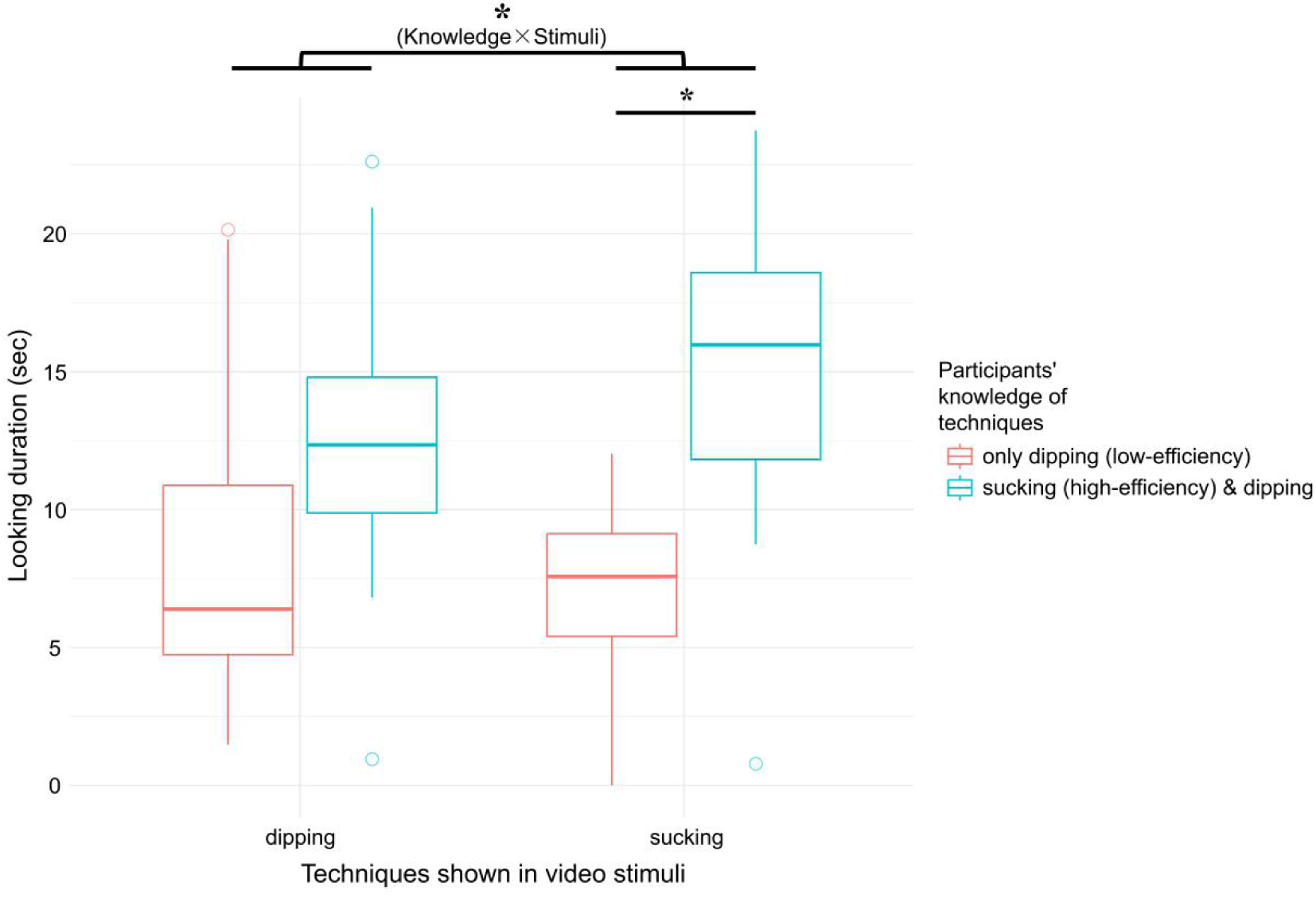
Looking duration to the screen by knowledge and stimuli. * *P* < 0.05. All participants know the dipping-technique, but only sucking-technique participants know the more efficient method, i.e. sucking.

The proportion of looking duration to each AOI is shown in Figure 3. Overall, both species looked similarly at the action AOI ( χ ^2^ = 0.868, *P* = 0.375), which remained consistent across trials ( χ^2^ = 0.0005, *P* = 0.983) and was not influenced by different demonstrators ( χ ^2^ = 0.496, *P* = 0.482), order of videos ( χ ^2^ = 0.0001, *P* = 0.994), tube-using knowledge ( χ ^2^ = 0.340, *P* = 0.574), or video stimuli ( χ ^2^ = 2.628, *P* = 0.116). For the face AOI, bonobos looked marginally longer compared to chimpanzees (β = - 0.192, SE = 0.093, χ^2^ = 4.237, *P* = 0.070). The trial number (χ^2^ = 3.074, *P* = 0.108), demonstrator (χ^2^ = 0.047, *P* = 0.828), video order (χ^2^ = 0.126, *P* = 0.723), tube-using knowledge (χ^2^ = 0.127, *P* = 0.730), and video stimuli (χ^2^ = 0.068, *P* = 0.799) did not have significant effects on participants’ attention to the face. For the food AOI, chimpanzees attended more than bonobos (β = 0.138, SE = 0.057, χ^2^ = 5.908, *P* = 0.036), while other predictors did not cause any differences (Trial: χ ^2^ = 0.252, *P* = 0.625, Demonstrator: χ^2^ = 0.593, *P* = 0.442, First-show: χ^2^ = 0.412, *P* = 0.522, Knowledge: χ^2^ = 0.303, *P* = 0.594, Stimuli: χ^2^ = 0.067, *P* = 0.799).

**Figure 3.**
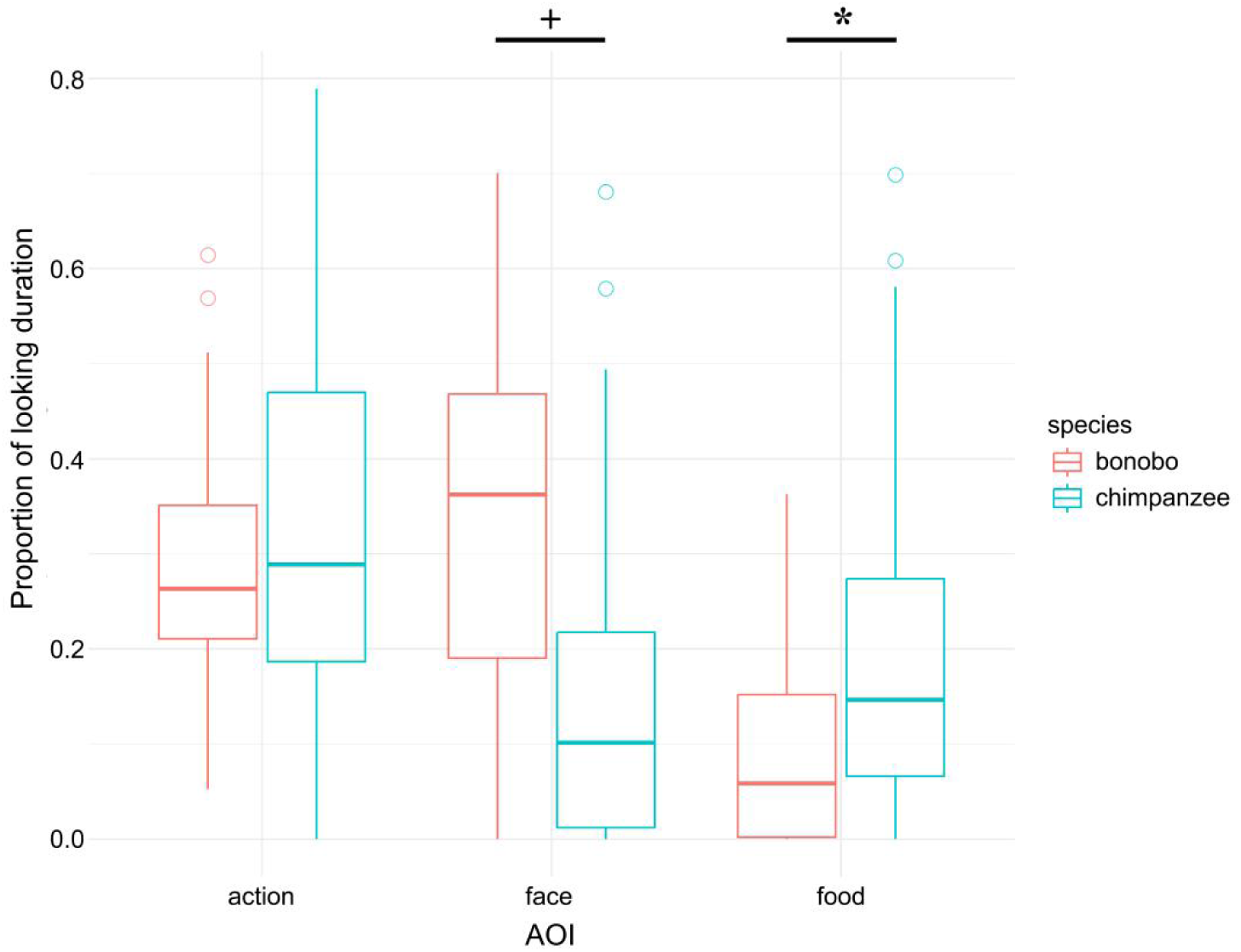
Proportion of looking duration to AOIs in bonobos and chimpanzees. + *P* < 0.1, * *P* < 0.05.

### Techniques employed following video demonstrations

Four of the six chimpanzees initially used the sucking technique while the other two used the dipping technique. All sucking-technique chimpanzees continued using this technique across the five trials, with the exception of one individual who on some trials did not attempt the task at all. No dipping-technique participants attempted to use the sucking technique following demonstrations, and both showed low overall motivation to use the tool. All six bonobos initially used the dipping technique, and none showed the sucking technique following video demonstrations. Surprisingly, only two individuals attempted to use the tool (dipping technique) in the test period, though others appeared interested in the juice by trying to use their fingers or tongue to reach it. Details and discussion of the coded behaviors of each participant can be found in supplementary materials.

## Discussion

This study tested how chimpanzees and bonobos viewed tool-using demonstrations that were same as/different from their known techniques through eye-tracking. We predicted that participants knowing different techniques would show different visual attention, and that learning success of chimpanzees and bonobos may differ. Results confirmed significant attentional differences to the screen and supported the first hypothesis, mainly driven by sucking-technique individuals looking significantly longer than dipping-technique individuals at sucking-technique videos. Thus, dipping-technique participants did not closely attend to demonstrations of a technique that they did not know. Both dipping- and sucking-technique participants paid similar attention to the action, face, and food part, indicating a similar attentional pattern to each AOI. As both species looked similarly at the action part, the second hypothesis was not fully supported, though a tendency for bonobos to attend more to faces was consistent with our prediction. None of the dipping-technique individuals of either species learned the sucking technique via the video demonstrations and our third hypothesis was therefore not supported.

While social learning from video demonstrations did not occur in our study, participants’ prior knowledge influenced their visual attention. Eye-tracking results showed that dipping-technique participants did not attend as closely to the unfamiliar sucking technique as those with prior knowledge of the technique. Close observation (peering) is a key factor for effective social learning [1,44,45], and as dipping-technique individuals did not closely attend to sucking videos it is therefore unsurprising they did not learn this technique from demonstrations. It should be noted that all participants are familiar with the dipping technique (as they use sticks to dip for juice during regular environmental enrichment), but only sucking-technique individuals knew the high-efficiency method, i.e. sucking. Due to the lack of prior knowledge [46], dipping-technique individuals only knew the low-efficiency method and likely did not understand the action of using the tube to suck. Because of the interplay of action observing, understanding, and execution [36,47,48], individuals that lack understanding of the action may additionally lack motivation to observe it.

Notably, apes often maintain their interest in easily understandable content while unfamiliar and complex content like animations and puppet plays generally fail to keep their attention [49]. Thus, apes may simply not pay much attention to what they do not know well. Given that the life-history of captive apes involves much opaque human tool-use, participants may be generally unmotivated to attend closely to human demonstrations. In this case, even familiar human demonstrators may not be suited for social learning of tool use. Low attention to human actions that are not (initially) understood as found here may partly explain failures to socially learn from humans in many past studies [50,51], despite abundant evidence for conspecific social learning [1,29,52]. It remains unclear whether using conspecific demonstrators might facilitate better attention and incur potential social learning in this paradigm. Future studies using eye-tracking with conspecific demonstrators could offer a more complete understanding of how they allocate their attention during the social learning process. Additionally, combined with aforementioned findings, for great apes, social learning of too-use may only occur in a limited zone [53,54]), where the ability to understand the action significantly influences the possibility of the individual observing and learning the behavior.

More broadly, several methodological factors may also suggest why dipping-technique participants failed to learn the sucking technique. Firstly, participants might not have received sufficient exposure to the demonstrations. In a previous study using the same task [33], dipping-technique chimpanzees underwent one to four 10-min trials with a conspecific demonstrator before adopting the sucking technique while our participants only received 48-second video demonstrations in each trial. However, as trials repeated, participants strongly decreased interest in the videos overall, suggesting that simply increasing exposure in this paradigm may not be sufficient. Second, non-human primates generally learn less effectively from video demonstrations compared with live demonstrations [52,55], though they can learn from videos in some contexts [56–58]. Two bonobos did seem to extract some information from video demonstrations (attempting to dip after ignoring the tool during familiarization trials in the testing room), though attention did not clearly differ in these trials and sample size was too small to draw any conclusions. Third, demonstrator identity may be central. Both human demonstrators were familiar to apes for many months or years, but participants did look less at the second demonstrator (Y.P.) who had known them for a shorter time. This may be due to an effect of demonstrator familiarity or may instead be due simply to participants quickly losing interest in repeated content. Future studies may be able to target each of these possibilities with more direct controlled conditions.

Our study used eye-tacking technology to investigate the social learning process of great apes, finding that apes may not closely observe things that they don’t know well (at least as performed by human demonstrators). This result offers potential explanations for some negative results in previous experiments and implications for the importance of using conspecific demonstrators in future studies. It also emphasizes the potential of applying eye-tracking to the study of social learning. With modified protocol and visual stimuli, such paradigms can also be used to study social learning in diverse species, providing insights into the foundational cognitive processes that underpin social learning and culture in non-human animals.

## Supplementary video

Video S1. Examples of eye-tracking recordings. Participants watching human experimenters (Y.P & J.B.) demonstrating the two techniques.

## Supporting information

Raw data

Supplemental information and details

## Acknowledgement

We sincerely appreciate the kind support from Dr. Naruki Morimura and Etsuko Nogami in conducting this study, as well as the great patience of apes at Kumamoto Sanctuary in participating in the study. We’d like to thank Ms. Yume Okamoto for her patient help in coding the recorded videos. This study was financially supported by Japan Society for Promotion of Science [JSPS KAKENHI 22KJ1884 to Y.P., 22KJ1677 to J.B., and 19H00629, 22H04451, and 24H02200 to S.Y.] and Japan

Science and Technology Agency [JST FOREST program JPMJFR221I to S.Y.].

## Ethics statement

All participants have much experience in joining in eye-tracking study and were tested in the usual testing rooms. Their daily participation in this study was voluntary. No changes were made to their daily care routine. Thus, they received regular feedings, daily enrichment, and had free access to water. This research was approved by the ethical committee of animal research in Wildlife Research Center, Kyoto University (WRC-2023-KS006A for chimpanzees, and WRC-2023-KS007A for bonobos).

## Declaration of interest

The authors declare no conflicts of interest.

## Authors’ contributions

Y.P.: methodology, investigation, formal analysis, writing - original draft, funding acquisition. J.B.: methodology, investigation, formal analysis, writing - review & editing, funding acquisition. S.Y.: conceptualization, project administration, writing - review & editing, funding acquisition, supervision.

## Notes

### Competing Interest Statement

The authors have declared no competing interest.

