## Supplemental information and details for "An eye-tracking study of visual attention in chimpanzees and bonobos when viewing different tool-using techniques"

**Methods**

1. Animals

Six chimpanzees (*Pan troglodytes verus*) and six bonobos (*Pan paniscus*) at Kumamoto Sanctuary, Japan, participated in this study (Table S1).

Table S1. Participant information

| Species | Age | Sex | Rearing history | Name | Knowledge |
| --- | --- | --- | --- | --- | --- |
| bonobo | 52 | F | Nursery-peer | Louise | dipping |
| bonobo | 33 | F | Nursery-peer | Ikela | dipping |
| bonobo | 35 | F | Nursery-peer | Lolita | dipping |
| bonobo | 42 | F | Mother | Lenore | dipping |
| bonobo | 29 | M | Nursery-peer | Junior | dipping |
| bonobo | 21 | M | Nursery-peer | Vijay | dipping |
| chimpanzee | 19 | F | Mother | Natsuki | sucking |
| chimpanzee | 28 | F | Nursery-peer | Mizuki | sucking |
| chimpanzee | 16 | F | Mother | Iroha | sucking |
| chimpanzee | 16 | F | Nursery-peer | Hatsuka | sucking |
| chimpanzee | 25 | F | Mother | Misaki | dipping |
| chimpanzee | 29 | M | Mother | Zamba | dipping |

Rearing history: “Mother” indicates individuals reared by their biological mothers, and “Nursery-peer” indicates individuals reared by human caretakers and conspecific peers.

Knowledge: “dipping” indicates individuals initially dipped the tube into the juice, and “sucking” indicates individuals initially used the tube to suck the juice.

*For more information of these apes, visit GAIN (https://shigen.nig.ac. jp/gain/, Great Ape Information Network: the online studbook of Japanese apes).

2. Location of the hole on the panel

After video demonstrations, participants could try the tube-using task themselves. A tube was provided through a hole on the panel, and the juice bottle was attached below this hole using a suction cup (diameter 25 mm). For chimpanzees, the hole (diameter 1.2 cm, height 55 cm above the ground) on the panel was the same one through which they sipped juice during eye-tracking. For bonobos, a new hole (diameter 1.8 cm, height 45 cm) was used because some participants tended to keep their mouth near the original hole and wait for the juice (functional fixedness).

3. Exploratory analyses

**Model 2:** **looking duration to eye and mouth** **AOI by Knowledge** **and Stimuli**

The dependent variables were the proportion of looking duration to eye and mouth AOI, i.e. the total looking duration to each AOI with respect to the total looking duration to the whole screen. And separate models were run for each dependent variable. The model syntax in R (similar to models reported in the main texts) was: the proportion of Looking duration ~ Trial + Species + Demonstrator + Knowledge*Stimuli + First-show + (1 + Trial + Stimuli || ID). We confirmed the normality of residuals, homogeneity of variance, and normality of random effects (ID) in both models and checked Variance Inflation Factors (VIF), which showed low collinearity (VIF < 5).

**Model 3: looking duration by Behavior and Stimuli**

To investigate whether participants’ behavioral responses in each tube-using task may indicate their motivation when viewing the video demonstrations, their behaviors in the task were video-coded and were classified into four types, (1) dipping: participants put one end of the tube into the juice and take it out, regardless of whether they lick that end; (2) sucking: participants use mouth to move the juice upwards inside the tube (higher than the liquid level), regardless of whether the juice reaches mouth; (3) quit: participants use the tube to touch the hole or insert through the hole, but without other following action; (4) none:participants do not take the tube or just throw it away. Participants’ behaviors in each tube-using task were shown in Table S2.

Total fixation duration was used for overall attention to the screen, and proportion of looking duration was used for each AOI, and separate models were run for each of these dependent variables. In these models, we included Trial (trial 1 to 5), Species (bonobos or chimpanzees), Demonstrator (first or second demonstrator), Behavior (dipping, sucking, quit, or none), Stimuli (dipping or sucking videos), the interaction between Behavior and Stimuli, and stimulus order (whether the first presented video was dipping/sucking), as fixed effects. We included participant identity (ID) to account for repeated measures for each individual, and the effects of Trial and Stimuli varying by ID, as random effects and random slopes, respectively. We confirmed the normality of residuals, homogeneity of variance, and normality of random effects (ID) in all models. We checked Variance Inflation Factors (VIF) through the “check_model”function and found that collinearity remained high for the interaction between behavior and stimuli in the models (VIF > 10). Therefore, we excluded the interaction of behavior and stimuli but still kept the two predictors. The modified model syntax was: the proportion of Looking duration ~ Trial + Species + Demonstrator + Behavior + Stimuli + First-show + (1 + Trial + Stimuli || ID). In these models, collinearity remained low (VIF < 5)

Table S2. Participants’ behaviors in the tube-using task of each trial

| Name | Species | Knowledge | Trial 1 | Trial 2 | Trial 3 | Trial 4 | Trial 5 |
| --- | --- | --- | --- | --- | --- | --- | --- |
| Louise | bonobo | dipping | none | none | none | none | none |
| Ikela | bonobo | dipping | none | quit | none | none | none |
| Lolita | bonobo | dipping | dipping | dipping | none | dipping | dipping |
| Lenore | bonobo | dipping | quit | none | none | none | none |
| Junior | bonobo | dipping | none | none | none | quit | none |
| Vijay | bonobo | dipping | none | dipping | quit | quit | dipping |
| Natsuki | chimpanzee | sucking | sucking | sucking | sucking | sucking | sucking |
| Mizuki | chimpanzee | sucking | sucking | sucking | sucking | sucking | sucking |
| Iroha | chimpanzee | sucking | sucking | sucking | sucking | sucking | sucking |
| Hatsuka | chimpanzee | sucking | sucking | quit | none | none | none |
| Misaki | chimpanzee | dipping | none | none | none | **quit** | **none** |
| Zamba | chimpanzee | dipping | none | none | **none** | **dipping** | **dipping** |

The results marked in bold indicates these trials were conducted in the room adjacent to the outdoor enclosure. See the above text for the description of participants’ behaviors.

**Results**

**Model 2: looking duration to eye and mouth AOI by Knowledge and Stimuli**

Looking duration to eye AOI

Participants looked rather similarly at demonstrators’ eye, which was not impacted by the main effects (Trial: χ^2^ = 0.461, *P* = 0.501; Species: χ^2^ = 0.677, *P* = 0.429; Demonstrator: χ^2^ = 0.076, *P* = 0.784; First-show: χ^2^ = 0.012, *P* = 0.914; Knowledge: χ^2^ = 0.000, *P* = 0.999; Stimuli: χ^2^ = 2.077, *P* = 0.164; the interaction between Knowledge and Stimuli: χ^2^ = 0.247, *P* = 0.628).

Looking duration to mouth AOI

Participants looked slightly less to the mouth AOI from Trial 1 to 5 (χ^2^ = 4.196, *P* = 0.066), and they looked marginally longer at sucking videos compared with dipping-technique videos (χ^2^ = 3.745, *P* = 0.065). But other main effects did not have significant influence on their visual attention to the demonstrators’ mouth (Species: χ^2^ = 1.796, *P* = 0.201; Demonstrator: χ^2^ = 0.043, *P* = 0.837; First-show: χ^2^ = 1.959, *P* = 0.167; Knowledge: χ^2^ = 1.488, *P* = 0.242; the interaction between Knowledge and Stimuli: χ^2^ = 0.362, *P* = 0.553).

**Model 3: looking duration by Behavior and Stimuli**

In the model using behavior as a fixed effect, Behavior was not a significant predictor of looking time for any dependent measure (whole screen and each AOI, *P* > 0.05).

Looking duration to whole screen

Participants’ looking duration to the whole screen significantly decreased across the five trials (χ^2^ = 22.001, *P* = 0.001) and they looked less at the second demonstrator of each video (χ^2^ = 25.039, *P* < 0.001). Other predictors did not cause any significant attentional differences (Species: χ^2^ = 0.766, *P* = 0.401; Behavior: χ^2^ = 0.972, *P* = 0.411; Stimuli: χ^2^ = 0.479, *P* = 0.503; First-show: χ^2^ = 0.468, *P* = 0.497).

Looking duration to action AOI

Chimpanzees looked marginally longer at the action AOI than bonobos (χ^2^ = 3.229, *P* = 0.098). But participants’ attention to action AOI was overall steady (Trial: χ^2^ = 0.017, *P* = 0.898; Demonstrator: χ^2^ = 0.499, *P* = 0.481; Behavior: χ^2^ = 1.029, *P* = 0.387; Stimuli: χ^2^ = 2.660, *P* = 0.115; First-show: χ^2^ = 0.009, *P* = 0.924).

Looking duration to face AOI

Chimpanzees looked slightly less at demonstrators’ face than bonobos (χ^2^ = 4.501, *P* = 0.058). But looking duration to the face AOI was not influenced by other predictors (Trial: χ^2^ = 2.597, *P* = 0.137; Demonstrator: χ^2^ = 0.049, *P* = 0.826; Behavior: χ^2^ = 1.679, *P* = 0.178; Stimuli: χ^2^ = 0.068, *P* = 0.799; First-show: χ^2^ = 0.589, *P* = 0.444).

Looking duration to food AOI

Chimpanzees looked significantly longer at the food AOI than bonobos (χ^2^ = 5.027, *P* = 0.046). But other predictors did not influence participants’ looking duration to the food AOI (Trial: χ^2^ = 0.288, *P* = 0.603; Demonstrator: χ^2^ = 0.590, *P* = 0.443; Behavior: χ^2^ = 0.290, *P* = 0.832; Stimuli: χ^2^ = 0.069, *P* = 0.796; First-show: χ^2^ = 0.594, *P* = 0.442).

**Techniques employed following video demonstrations**

All sucking-technique chimpanzees continued using this technique across the five trials, with the exception of a nursery-peer chimpanzee, Hatsuka. This individual seemed to want human attention more than the grape juice and kept trying to demand for something else. Therefore, she did not attempt the task at all in some trials. The two dipping-technique participants were often eager to leave the testing room and reluctant to use the plastic tube after the first several trials. When tested in the room connected to the outdoor enclosure, Zamba started to use the tube as a dipping tool, while Misaki was still not interested and only inserted the tube into the juice bottle and left in Trial 4.

All six bonobos initially used the dipping technique. However, four of them stopped using the provided tubes in the five trials, though they still showed some interest in the grape juice by trying to use their fingers or tongue to reach the juice through the hole, especially Ikela, Lenore, and Junior. The other two participants, Lolita and Vijay, still accepted the plastic tube and dipped it into the juice.

Before data collection, participants underwent several familiarization trials using unrelated videos (mating animals and social play), which familiarized them with the procedure of joining in the eye-tracking first and then staying a while for the task. During this phase, all chimpanzees continued using their own techniques while all bonobos stopped taking the tubes and tried to use their fingers and tongue instead. When real trials started, Lolita immediately recovered her dipping technique in Trial 1 and Vijay did so in Trial 2. Thus, the video demonstrations might still remind these two bonobos of their technique, though their attention did not show any differences and this sample size was rather limited.

During regular environmental enrichment, all participants often use natural sticks to dip the juice. Because the provided tubes were much less efficient for dipping than the sticks they use, the lack of use by participants may have reflected dissatisfaction with the tool for dipping rather than lack of interest in the juice per se, especially some participants did try to reach the juice using their fingers or tongue.
